# The development of a novel diagnostic PCR for *Madurella mycetomatis* using a comparative genome approach

**DOI:** 10.1101/2020.04.10.036681

**Authors:** Wilson Lim, Kimberly Eadie, Emmanuel Siddig, Bertrand Nyuykonge, Sarah Ahmed, Ahmed H. Fahal, Annelies Verbon, Sandra Smit, Wendy WJ van de Sande

## Abstract

Eumycetoma is a neglected tropical disease characterized by large tumorous lesions. It is most commonly caused by the fungus *Madurella mycetomatis* which accounts for more than 70% of cases in central Africa. Currently, identification of the causative agent can only be reliably performed by a species-specific PCR. However, we recently demonstrated that our *M. mycetomatis* specific PCR can cross-react with *Madurella pseudomycetomatis.* We therefore used a comparative genome approach to develop a new *M. mycetomatis* specific PCR for species identification. For this we compared the published *M. mycetomatis* genome to genomes of other organisms in BLASTCLUST to identify unique *M. mycetomatis* predicted protein coding sequences. Based on 16 of these unique sequences, PCR primers were developed. The specificity of these primers was further evaluated in other eumycetoma causing agents including the *Madurella* sibling species. Out of the 16 tested sequences, only one was unique for *M. mycetomatis* and this should be used as a novel diagnostic marker for *M. mycetomatis.*

## INTRODUCTION

The neglected tropical disease mycetoma presents itself as a subcutaneous chronic granulomatous infectious and inflammatory disease and is characterized by tumorous lesions (1, 2). This disease can be caused by more than 70 different micro-organisms and is categorized into actinomycetoma (caused by bacteria) and eumycetoma (caused by fungi). Most cases occur in the mycetoma belt between the latitudes 15° South and 30° North. Diagnosis of eumycetoma is often only made clinically in endemic areas due to the scarcity of facilities, expertise and financial capacity. Identification of the causative agent is time consuming and often limited to culture and histology which can leads to misidentifications (1, 3, 4). The only way to properly identify eumycetoma causative agents to the species level is through molecular identification, commonly PCR. PCR identification’s reliability, high turnover and sensitivity has made it widely used in diagnosing fungal disease, and in most cases, it has already replaced culturing of the microorganism as the primary diagnostic method (5-11).

*Madurella mycetomatis* is a fungus only recognised as a pathogen in mycetoma and is responsible for more than 70% of all mycetoma infections in endemic areas (1, 12, 13). In 1999, specific PCR primers based on the internal transcribed spacer (ITS) region were developed for *M. mycetomatis* by our group (14), however, recently, this *M. mycetomatis* specific PCR primer pair was discovered to cross-react with *Madurella pseudomycetomatis* (15). Back then, *M. pseudomycetomatis* was not yet discovered (16). Since new fungi causing eumycetoma are still being discovered, there is clearly a need for a specific PCR marker to identify *M. mycetomatis. M. pseudomycetomatis* are more commonly found in Central and South America, while *M. mycetomatis* is predominant in the African continent (17). All four *Madurella* species (*Madurella fahalii, Madurella tropicana, M. mycetomatis* and *M. pseudomycetomatis*) are known to cause mycetoma and requires different treatment strategies, thus, it is important to be able to distinguish between them in order to study their epidemiology and to administer proper treatment (18). Since all four *Madurella* species share a very conserved ITS region, this has made designing PCR primers specific for *M. mycetomatis* based on that region difficult (18, 19). To avoid the complication brought by the ITS region, we took a different approach to design specific primers to diagnose *M. mycetomatis* using PCR. Here, using the unique conserved DNA sequence from the published genome of *M. mycetomatis* (20), we have identified diagnostic DNA markers and developed a new species specific PCR to diagnose *M. mycetomatis.*

## MATERIALS AND METHOD

### Fungal isolates

A total of 93 fungal isolates were used in this study; 60 *M. mycetomatis*, 4 *M. tropicana,* 3 *M. fahalii,* 3 *M. pseudomycetomatis,* 1 *Aspergillus fumigatus,* 1 *Aspergillus terreus,* 2 *Chaetomium globosum,* 4 *Falciformispora senegalensis,* 1 *Fusarium solani,* 3 *Medicopsis romeroi,* 3 *Thielavia terrestris,* 3 *Thielavia subthermophilia,* 4 *Trematospheria grisea* and 1 *Trichophyton rubrum.* These fungal isolates were obtained from both the Mycetoma Research Center in Sudan and the Westerdijk Fungal Biodiversity Institute in the Netherlands and maintained in Erasmus Medical Centre. All isolates were identified to the species level on the basis of morphology, polymerase chain reaction (PCR)-based restriction fragment length polymorphisms, and sequencing of the ITS regions (3, 14, 21).

### DNA isolation

Fungal isolates were first grown on Sabouraud Dextrose agar (Difco Laboratories) with or without gentamicin (40mg/mL, Centrafarm Nederland B.V., The Netherlands) for three weeks at either 37°C or room temperature depending on the fungal species. The mycelium was then harvested from the agar and subjected to 3 minutes of bead bashing (Tissuelyser, Qiagen) at maximum power of 30 frequency per second using ten to twelve 3mm metal beads (DIT Holland B.V., The Netherlands) per 2mL Eppendorf tube. DNA was then isolated using ZR Fungal/Bacterial DNA MicroPrepTM kit (Zymo Research, Irvine, California, USA) according to manufacturer instructions while omitting the bead bashing step from the kit.

### Identifying unique predicted protein coding sequences in the *M. mycetomatis* genome

*M. mycetomatis* predicted protein coding sequences were obtained from the recently published genome sequence of *M. mycetomatis* isolate mm55, accession number LCTW00000000, BioProject PRJNA267680 (20). To determine their specificity to *M. mycetomatis*, a bioinformatical comparison between the predicted protein coding sequences and the genome of other organisms was performed using BLASTCLUST (22). Predicted protein coding sequences were chosen depending on their orthologues, E-value and fragment size. Orthologues were defined as predicted amino acid sequences with greater than 85% amino acid identity to *M. mycetomatis* proteins. Predicted protein coding sequences with no orthologues present in the genomes of other organisms was chosen based on the E-value determined through BLASTCLUST. Predicted protein coding sequences with an E-value of 0.003 and higher was selected as a cut-off point and considered to be specific to *M. mycetomatis* and those between 400 bp and 1100 bp was further selected for analysis.

### Primer design and PCR conditions

Forward and reverse primers were designed according to the nucleotide sequence of the predicted protein coding sequences of interest. Primer sequences are depicted in Table 1. PCR reaction was set up with to a final volume of 25μl containing 0.6 units of Super Taq HC DNA polymerase (Sphaero Q), 0.1 nM/μl DNTP (Thermo Fisher Scientific) and 0.5 pmol/μl of each forward and reverse primer. DNA was amplified in a thermal cycler (Applied Biosystems VeritiTM) using the following program: initial denaturation at 94°C for 10 min; 40 cycles of amplification with various annealing temperatures (95°C for 1 minute, 55-59°C for 1 minute, and 72°C for 1 minute); and a final extension step of 10 seconds at 72°C. The PCR reaction products were visualized using 2% agarose gel (Sphaero Q) with GeneRuler 100bp Plus DNA ladder (Thermo Fisher Scientific) and stained with SYBR^®^ Safe DNA Gel Stain (Thermo Fisher Scientific). Bands on gel were visualized using a gel imaging machine (Isogen Life Science B. V., The Netherlands).

**Table 1.**
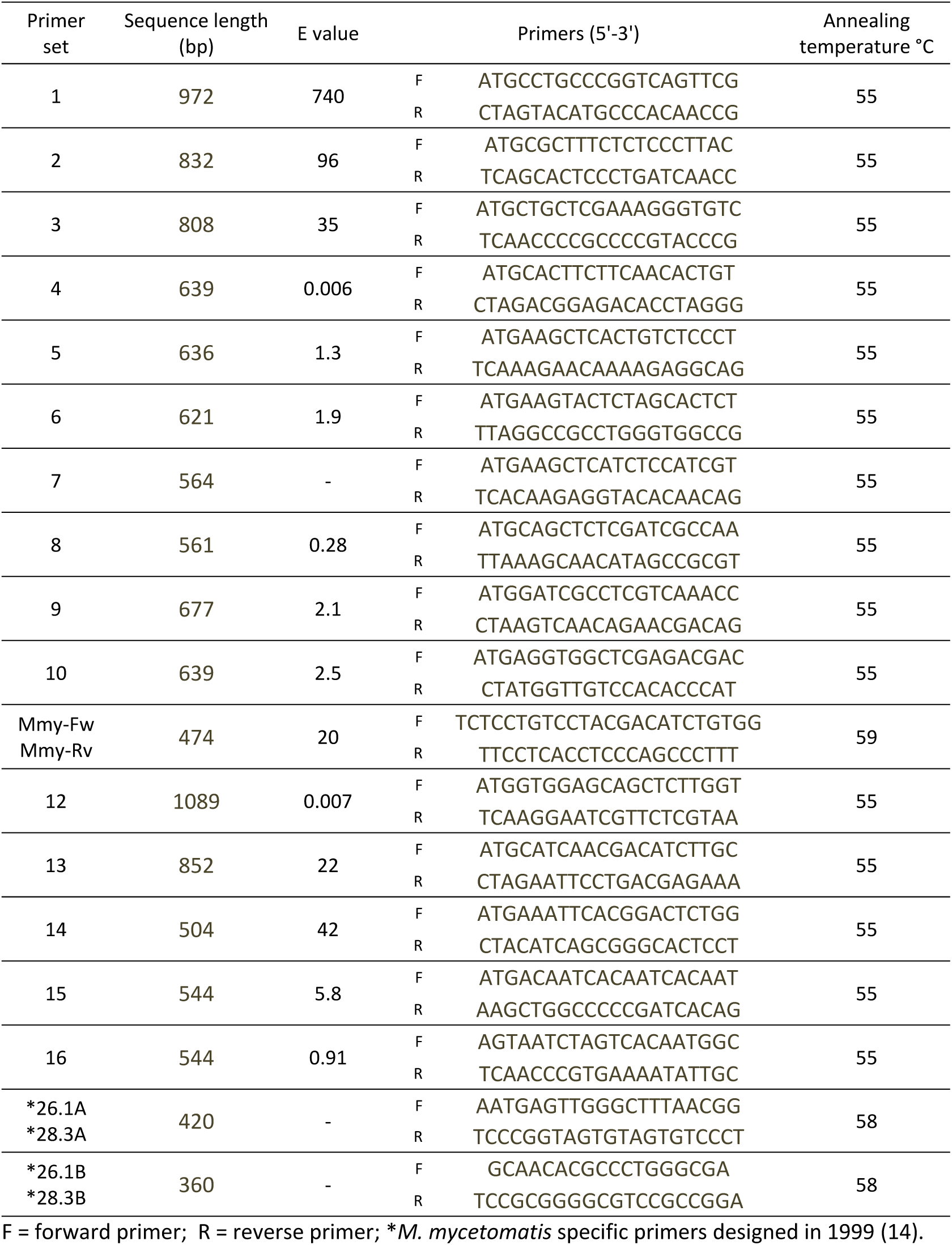
The sixteen predicted protein sequences with their corresponding size, primer sequences and annealing temperatures.

## RESULTS and DISCUSSION

Recently, the currently used *M. mycetomatis* specific PCR primers were discovered to cross-react with *M. pseudomycetomatis* (15). To be able to study the epidemiology of the fungi and to administer proper treatment on patients, it is important to be able to distinguish between the different fungal species that cause eumycetoma, therefore, there is a need to design *M. mycetomatis* specific primers for diagnostic purposes. From the genome of *M. mycetomatis,* a total of 350 predicted protein coding sequences were selected and analyzed. We chose to select predicted protein coding sequences because protein coding regions are likely to be more stable than non-coding (23, 24). To ensure that they can be easily amplified through PCR, we preferentially chose predicted protein coding sequences with sizes between 400 and 1100bp. From the initial 350 predicted protein coding sequences, the top 16 candidates that fitted our requirement were chosen for PCR development.

PCR primers for the 16 candidates were then designed (Table 1). To ascertain that the primers would amplify their targets in all *M. mycetomatis* isolates, the PCR primers were evaluated in 60 *M. mycetomatis* isolates from different geographical origin, genotypic background and phenotypic appearance. Out of the 16 primer sets tested, 13 were positive in all *M. mycetomatis* isolates (figure 1). Primer sets 4, 5 and 12 are present in 58, 4 and 59 isolates, respectively (figure 1). To determine the specificity of the 13 positive primer sets, they were tested against other fungal mycetoma causative agents and close relatives of *M. mycetomatis*. As seen in table 2, only primer set 11 – later renamed as Mmy-Fw and Mmy-Rv - was found to be specific for *M. mycetomatis*. Primer set 2, 4, 8 and 9 were not able to discriminate between the different *Madurella* species while 5 and 7 could discriminate between the four *Madurella* species but cross reacted with at least one other mycetoma causative agent. Mmy-Fw and Mmy-Rv appears to be a putative single-copy gene, making it an ideal candidate as an identification marker. Since it seemed the most suited for identifying *M. mycetomatis,* it was compared to the currently used diagnostic PCR to determine the limit of detection. Mmy-Fw and Mmy-Rv was observed to be only slightly less sensitive compared to the current *M. mycetomatis* specific PCR primer pair 26.1a and 28.3a. Mmy-Fw and Mmy-Rv was able to detect DNA concentrations as low as 5 pg whereas the slightly more sensitive *M. mycetomatis* specific PCR was able to detect DNA at 0.5 pg.

**Table 2.** Presence or absence of PCR amplicons of the sixteen primer sets and PCR primers developed in 1999 (14) in the other eumycetoma causing agents and close relatives of *M. mycetomatis.* The grey box highlights the absence of amplicons in all species tested here using Mmy-Fw and Mmy-Rv. Only PCR with bands of the same sizes to *M. mycetomatis* are considered specific to *M. mycetomatis*.

| Primer set | 1 | 2 | 3 | 4 | 5 | 6 | 7 | 8 | 9 | 10 | Mmy-Fw<br>Mmy-Rv | 12 | 13 | 14 | 15 | 16 | *<br>26.1A<br>28.3A | *<br>26.1B<br>28.3B |
| --- | --- | --- | --- | --- | --- | --- | --- | --- | --- | --- | --- | --- | --- | --- | --- | --- | --- | --- |
| *Madurella Tropicana* (4) | A | A | C | A | C | A | C | A | A | A | C | A | C | B | B | C | C | A |
| *Madurella fahalii* (3) | C | A | C | A | C | C | C | A | A | A | C | C | C | B | B | B | C | A |
| *Madurella pseudomycetomatis* (3) | A | A | A | A | C | B | C | A | A | B | C | B | B | C | B | A | B | A |
| *Aspergillus fumigatus* (1) | - | - | - | - | - | - | C | - | - | - | C | - | - | - | - | - | - | - |
| *Aspergillus terreus* (1) | - | - | - | - | - | - | C | - | - | - | C | - | - | - | - | - | - | - |
| *Chaetomium globosum* (2) | - | - | - | - | - | - | B | - | - | - | C | - | - | - | - | - | - | - |
| *Falciformispora senegalensis* (4) | B | B | B | B | B | B | C | B | B | B | C | C | B | B | C | B | C | C |
| *Fusarium solani* (1) | C | C | B | C | B | B | C | B | B | A | C | C | B | B | B | B | C | C |
| *Medicopsis romeroi* (3) | - | - | - | - | - | - | A | - | - | - | C | - | - | - | - | - | C | C |
| *Thielavia subthermophila* (3) | B | C | B | C | C | B | C | B | B | B | C | B | B | B | B | C | C | C |
| *Thielavia terrestris* (3) | B | B | B | C | C | B | C | B | B | B | C | B | B | B | B | C | C | C |
| *Trematospheria grisea* (4) | - | - | - | - | - | - | C | - | - | - | C | - | - | - | - | - | C | C |
| *Trichophyton rubrum* (1) | - | - | - | - | - | - | C | - | - | - | C | - | - | - | - | - | C | C |

A: PCR band of the same size; B: PCR band of another size; C: no PCR band. \**M. mycetomatis* specific primers designed in 1999 (14).

**Figure 1.**
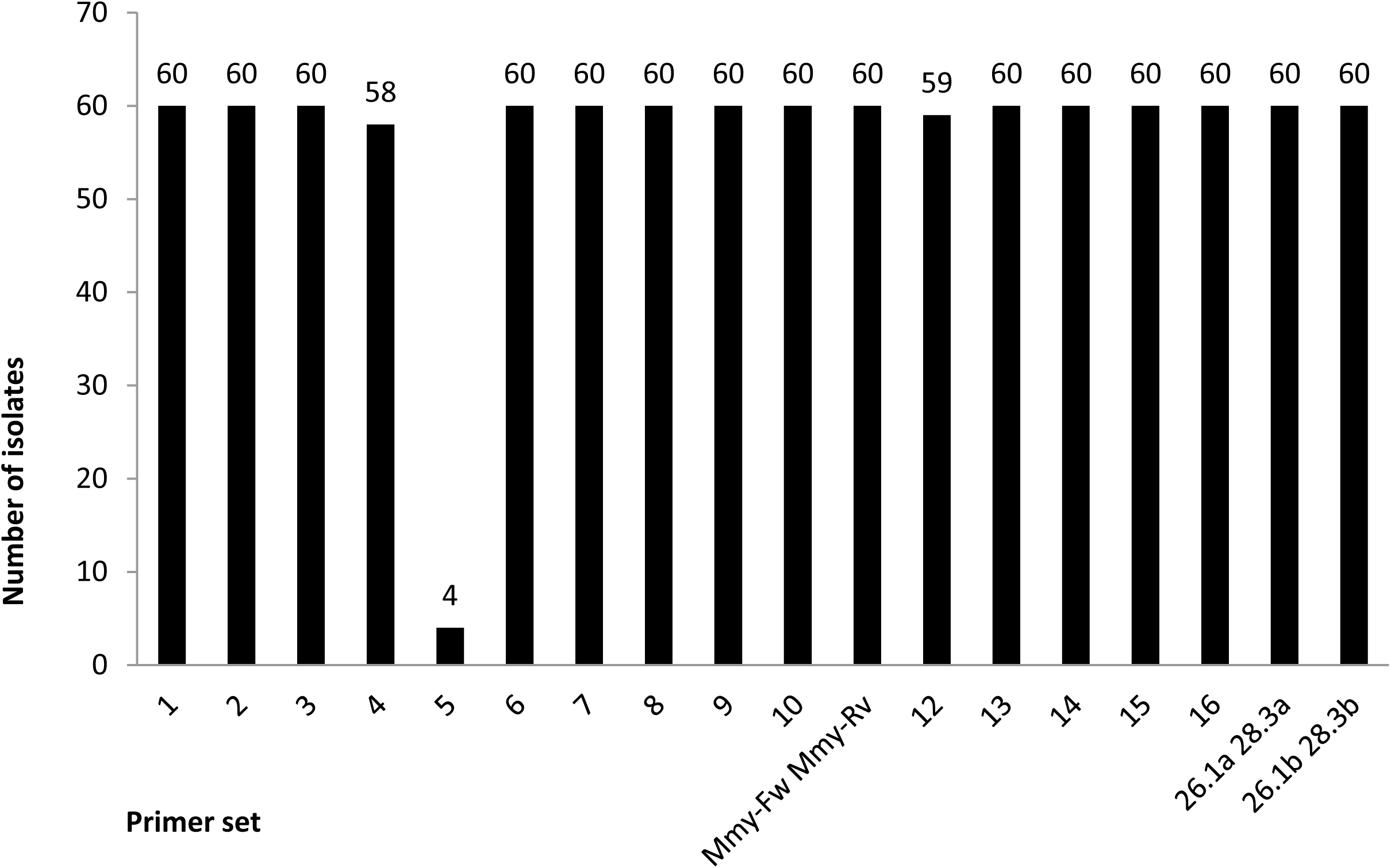
Presence of the 16 PCR amplicons in 60 *M. mycetomatis* isolates tested. Most PCR reactions resulted in amplification in all isolates tested except PCR 4, 5 and 12.

One of the advantages of the comparative genome method is that primer design is less constrained since the targeted genes are unique. With this method, we were able to design primers that are able to distinguish between *M. mycetomatis* and *M. pseudomycetomatis.* Using this approach, other studies also succeeded in designing specific primers for their organism of choice (25-27). In a study by Withers *et al*, a similar genome comparison method was performed on *Pseudoperonospora cubensis* and *Pseudoperonospora humuli* (27). The comparison was first performed *in silico* and subsequently *in vitro.* Using this approach, they were able to identify and determine a large number of specific markers for their organism of interest (27). However, we were not able to perform a similar *in silico* approach here because at the time of data analysis and the preparation of this manuscript, only one *M. mycetomatis* isolate has ever been sequenced and none of *M. fahalii, M. tropicana* and *M. pseudomycetomatis* has ever been sequenced.

In conclusion, since cross-reactivity occurs with the current *M. mycetomatis* specific PCR primer pair 26.1a and 28.3a, we have used a comparative genome approach to identify and designed a new *M. mycetomatis* species-specific PCR primers. We now recommend all reference and local laboratories to use the new PCR primers Mmy-Fw and Mmy-Rv to identify *M. mycetomatis* to the species level. Furthermore, this comparative genome approach may also be used to design markers for other eumycetoma agents and also other fungi that share conserve ITS region within its genus.

